# Argininosuccinic aciduria fosters neuronal nitrosative stress reversed by *Asl* gene transfer

**DOI:** 10.1101/348292

**Authors:** Julien Baruteau, Dany P. Perocheau, Joanna Hanley, Eridan Rocha-Ferreira, Rajvinder Karda, Joanne Ng, Nattalie Suff, Ahad A. Rahim, Michael P. Hughes, Blerida Banushi, Helen Prunty, Mariya Hristova, Deborah A. Ridout, Alex Virasami, Simon Heales, Stewen J. Howe, Suzy M. Buckley, Philippa B. Mills, Paul Gissen, Simon N. Waddington

## Abstract

Argininosuccinate lyase (ASL) belongs to the liver-based urea cycle detoxifying ammonia, and the citrulline-nitric oxide cycle synthesising nitric oxide (NO). ASL-deficient patients present argininosuccinic aciduria characterised by hyperammonaemia and a multi-organ disease with neurocognitive impairment. Current therapeutic guidelines aim to control ammonaemia without considering the systemic NO imbalance. Here, we observed a neuronal disease with oxidative/nitrosative stress in ASL-deficient mouse brains. A single systemic injection of gene therapy mediated by an adeno-associated viral vector serotype 8 (AAV8) in adult or neonatal mice demonstrated the long-term correction of the urea cycle and the citrulline-NO cycle in the brain, respectively. The neuronal disease persisted if ammonaemia only was normalised but was dramatically reduced after correction of both ammonaemia and neuronal ASL activity. This was correlated with behavioural improvement and a decrease of the cortical cell death rate. Thus, the cerebral disease in argininosuccinic aciduria involves neuronal oxidative/nitrosative stress not mediated by hyperammonaemia, which is reversed by AAV gene transfer targeting the brain and the liver, acting on two different metabolic pathways *via* a single vector delivered systemically. This approach provides new hope for hepatocerebral metabolic diseases.

## MAIN TEXT

Adeno-associated virus (AAV) vector mediated gene therapy has achieved promising results in recent clinical trials in liver ^1^ and neurodegenerative ^2^ inherited diseases, and led to the market approval of the first gene therapy product in the western world to treat lipoprotein lipase deficiency ^3^. These successes underpin the current interest in this technology as illustrated by the rapidly expanding number of gene therapy based clinical trials ^4^. Among various AAV capsid variants, AAV8 has demonstrated its efficacy in liver transduction in preclinical ^5^ and clinical studies ^6^. This serotype also efficiently transduces other tissues including the central nervous system after systemic injection in neonatal mice ^7^.

As with many liver inherited metabolic diseases, urea cycle defects exhibit a high rate of mortality and neurological morbidity in infancy despite conventional treatment ^8^. Successful correction of the urea cycle *via* AAV-mediated gene therapy has been reported in mouse models of ornithine transcarbamylase deficiency ^9,10^, argininosuccinate synthetase deficiency ^11,12^, and arginase deficiency ^13^. Argininosuccinic aciduria (ASA; OMIM 207900) is the second most common urea cycle defect with a prevalence of 1/218,000 live births ^14^ In addition, ASA is an inherited condition proven to cause systemic nitric oxide (NO) deficiency ^15^ as the disease is caused by mutations in argininosuccinate lyase (ASL), an enzyme involved in two metabolic pathways: i) the liver based urea cycle which detoxifies ammonia, a highly neurotoxic compound generated by protein catabolism and ii) the citrulline-NO cycle, present in most organs, producing NO from L-arginine *via* nitric oxide synthase (NOS) (Supplementary Fig. 1) ^16^. Patients may exhibit an early-onset phenotype with hyperammonaemic coma in the first 28 days of life, or a late-onset phenotype with either acute hyperammonaemia or a chronic phenotype with neurocognitive impairment and progressive liver disease ^17^. Compared to other urea cycle defects, ASA patients present with an unusually high rate of neurological and systemic complications ^17^ contrasting with a lower rate of hyperammonaemic episodes. Various pathophysiological mechanisms have been hypothesised to account for this paradox, including impaired NO metabolism ^18^. A hypomorphic *Asl*^*Neo/Neo*^ mouse model shows impairment of both urea and citrulline-NO cycles and reproduces the clinical phenotype with impaired growth, multi-organ disease, hyperammonaemia and early death ^15^. Common biomarkers of ASA include increased ammonaemia, citrullinaemia, plasma argininosuccinic acid, orotic aciduria and reduced argininaemia ^18^.

In this study, we have characterised the neuropathophysiology of the disease studying the brain of the hyperammonaemic *Asl*^*Neo/Neo*^ mouse and have used a systemic AAV-mediated gene therapy approach as a proof-of-concept study to rescue survival and protect the ASL-deficient brain from both hyperammonaemia and cerebral impaired NO metabolism. To achieve this, we designed a single-stranded AAV8 vector carrying the murine *Asl* (*mAsl*) gene under transcriptional control of a ubiquitous promoter, the short version of the elongation factor 1 α (EFS) promoter. The vector was administered systemically to adult and neonatal *Asl*^*Neo/Neo*^ mouse cohorts.

## RESULTS

### Pathophysiology of the brain disease in ASA

ASL deficiency causes a systemic NO deficiency due to the loss of a protein complex that facilitates channeling of exogenous L-arginine to NOS ^15^. To explore the effect on cerebral NO metabolism, various surrogate biomarkers were investigated. NO concentrations from wild type (WT) and *Asl*^*Neo/Neo*^ mice were evaluated by measurement of nitrite (NO_2_^−^) and nitrate (NO_3_^−^) ions, downstream metabolites of NO ^19^, and were found to be significantly increased in *Asl*^*Neo/Neo*^ mice in brain homogenates (Fig. 1A). Similarly, cyclic guanosine monophosphate (cGMP), .a signalling pathway physiologically upregulated by NO generated by coupled NOS ^20^, when measured in brain homogenates, was also found to be increased in *Asl*^*Neo/Neo*^ mice (Fig. 1B). Low tissue L-arginine is a consequence of ASL deficiency downstream the metabolic block and can cause NOS uncoupling ^21^, which leads to the production of reactive oxygen species including superoxide ion (O_2_^−^) or peroxynitrite (ONOO^−^) with the latter nitrating specific tyrosine residues and generating nitrotyrosine, a marker of oxidative/nitrosative stress ^22^. This process can modify the protein structure and function, altering enzymatic activity or triggering immune response ^23^. The detoxification of peroxynitrite by reduced glutathione (GSH) can generate nitrite *via* the reaction ONOO^−^ + 2GSH → NO_2_^−^ + GSSG + H_2_O ^24^. Analysis of glutathione concentrations in in brain homogenates of *Asl*^*Neo/Neo*^ mice showed that they were reduced although this did not reach statistical significance (Fig. 1C). In the cortex of WT and *Asl*^*Neo/Neo*^ mice, immunostaining against nitrotyrosine was increased in *Asl*^*Neo/Neo*^ mice in cells identified morphologically as neurons (Fig. 1D). Staining of glial fibrillary acidic protein (GFAP) and CD68, markers of astrocytic and microglial activation, respectively, did not show any difference (Fig. 1E and 1F, respectively). Immunohistochemistry against neuronal NOS (nNOS or NOS1), inducible NOS (iNOS or NOS2) and endothelial NOS (eNOS or NOS3) showed an increased staining of all three enzymes in *Asl*^*Neo/Neo*^ mice in cells morphologically identified as neurons (nNOS and iNOS) and endothelial cells (eNOS) (Fig. 1G, 1H and 1I, respectively). The brain morphology did not differ between WT and *Asl*^*Neo/Neo*^ mice (Supplementary Fig. 2). Collectively these data suggest that a neuronal oxidative/nitrosative stress plays a role in the neuropathology of ASA. However hyperammonaemia *per se* can cause brain toxicity through oxidative stress ^25^. To investigate whether neuronal oxidative/nitrosative stress is a primary mechanism involved in the phenotype of patients with ASA or is secondary to hyperammonaemia, we designed a gene therapy approach to normalise ammonaemia and conditionally target neuronal ASL activity.

**Figure 1.**
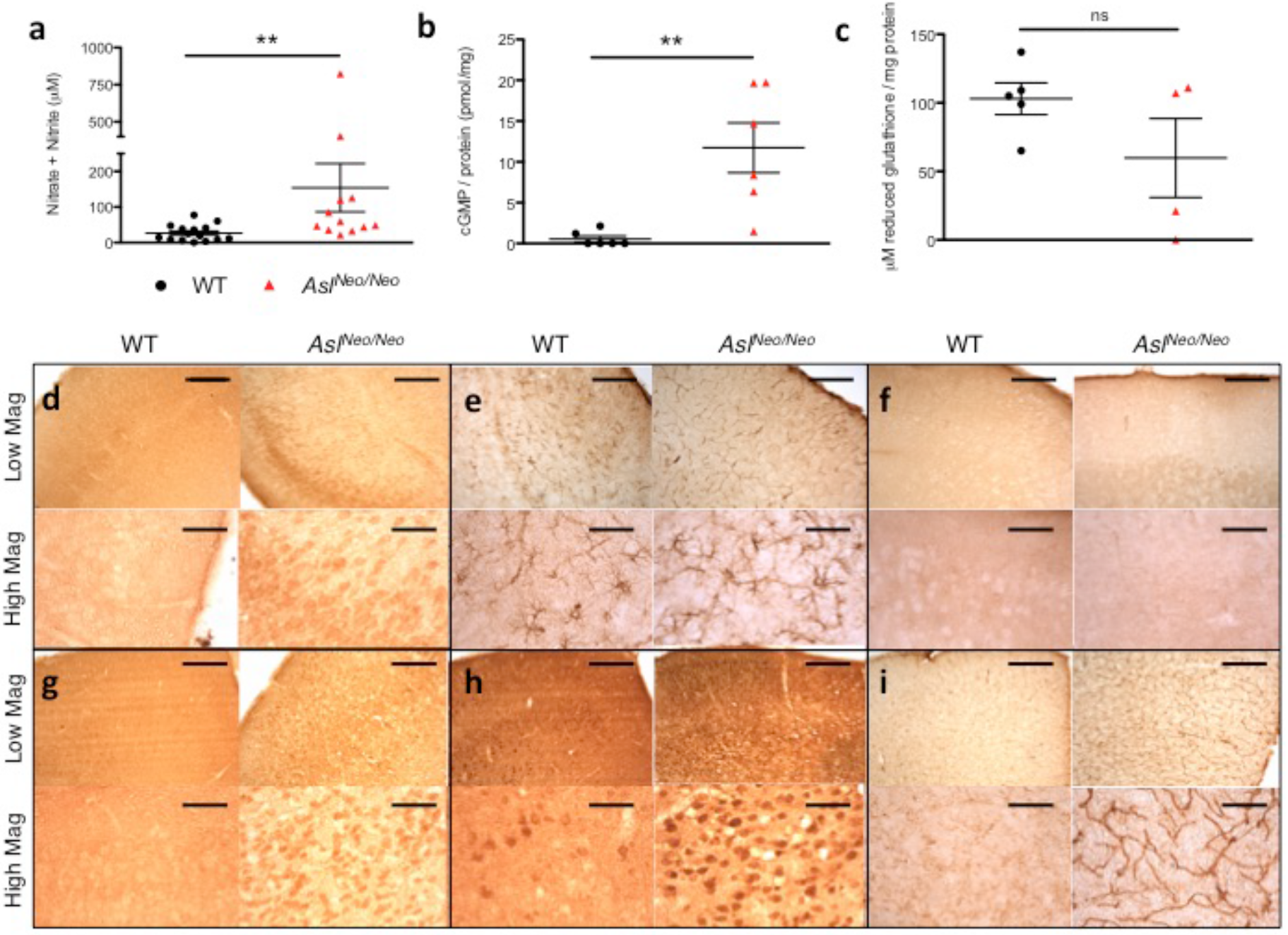
Neuronal oxidative/nitrosative stress is a component of the neurological disease in *Asl*^*Neo/Neo*^ mice. **(a)** Nitrite/nitrate levels (n=12-15) **(b)** cyclic GMP (n=6) and **(c)** reduced glutathione (n=4-5) in brain homogenates. Immunostaining of cortical sections shows **(d)** increased nitrotyrosine in morphologically-identified neurons, **(e)** no difference in the astrocytic GFAP or **(f)** microglial CD68, but increased nNOS **(g)** and iNOS **(h)** in morphologically-identified neurons and **(i)** eNOS in endothelial cells of 1-3 month-old WT and *Asl*^*Neo/Neo*^ mice (n=3). Horizontal lines display the mean ± standard error of the mean (SEM). ns = not significant. Unpaired 2-tailed Student’s *t* test ** p<0.01. **(a)** Graph displays not transformed data. Log transformed data were used for statistical analysis. Scale bars of low and high magnification: 500 and 125 μm, respectively.

### AAV8.EFS.GFP vector targets liver and cerebral neurons

In order to extend survival and ameliorate the brain phenotype, we designed a vector that was not only able to transduce the liver to correct the defective urea cycle but also the brain, especially neurons. Neonatal C57Bl/6 mice received an intravenous injection of a single stranded AAV8.EFS.*GFP* vector (3.4×10^11^vector genomes/pup) and were culled at 5 weeks of life alongside uninjected control littermates. Fluorescence microscopy revealed green fluorescent protein (GFP) expression in the liver and the brain (Fig. 2A). Anti-GFP immunostaining confirmed a high rate of hepatocyte transduction across the hepatic lobule (Fig. 2B). Anti-GFP ELISA showed the liver as the main peripheral organ transduced (Supplementary Fig. 3A, 3B). Anti-GFP brain immunostaining showed a clear pattern of neuronal transduction (Fig. 2C, 2D) throughout, most prominently in the cortex and decreasing rostro-caudally (Supplementary Fig. 3C).

**Figure 2.**
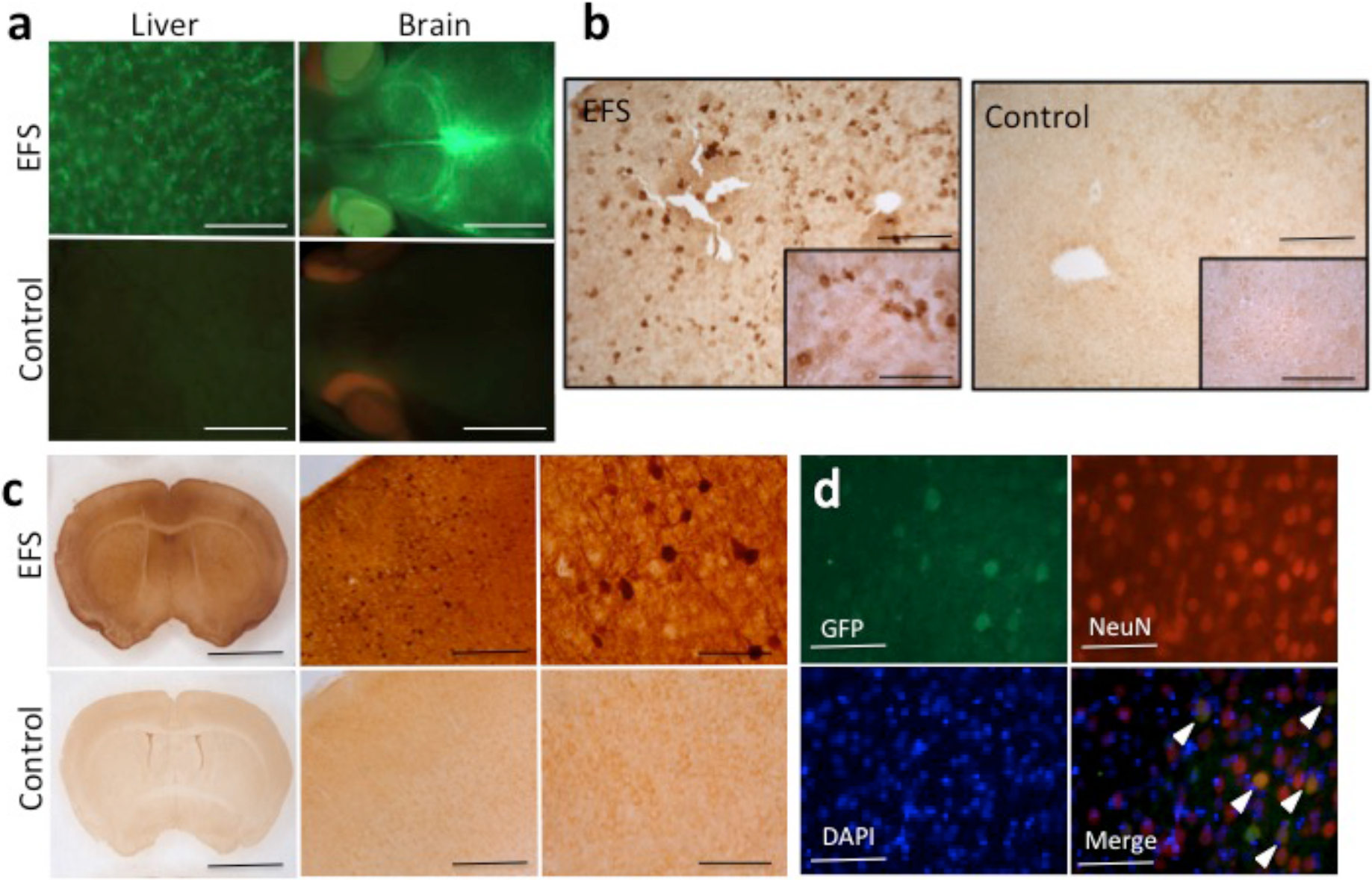
Neonatal intravenous injection of AAV8.EFS.GFP shows liver and brain transduction. **(a)** Liver and brain imaged with fluorescence microscope in mice injected with AAV8.EFS.*GFP* (AAV) and uninjected controls. Representative images of GFP immunostaining in liver **(b)** and in brain **(c)** at higher magnifications in mice injected with the EFS vector and uninjected controls. **(d)** Immunofluorescence of cortical staining for GFP (green), NeuN (red) and DAPI (blue). Arrows identify neurons. Representative images obtained by confocal microscopy in mice injected with the AAV vector; scale bars: 5 mm **(a)**; 500 μm and 125 μm in main and inset pictures, respectively **(b)**; from left to right 5 mm, 500 μm and 125 μm **(c)**; 500 μm **(d)**; n=4.

### Long-term correction of the macroscopic phenotype of Asl^Neo/Neo^ mice after adult but not neonatal injection of AAV8 gene therapy

*Asl*^*Neo/Neo*^ mice received supportive treatment (L-arginine and sodium benzoate, see Methods) allowing improved survival until day 30 (considered as adulthood; Supplementary Fig. 4A). A group of *Asl*^*Neo/Neo*^ mice received a single intraperitoneal injection of AAV8.EFS*.mAsl* (2.5×10^11^vg/mouse). Supportive treatment was withdrawn the following day for all *Asl*^*Neo/Neo*^ mice that had received gene therapy (adult-injected *Asl*^*Neo/Neo*^ mice) or not (untreated *Asl*^*Neo/Neo*^ mice). The mice were monitored until sacrifice at 12 months after injection. Another group of *Asl*^*Neo/Neo*^ (neonatal) mice were injected within 24 hours of birth with a single intravenous injection of AAV8.EFS*.mAsl* (3.2×10^11^vg/mouse). The neonatally-injected mice did not receive any supportive treatment and were monitored until sacrifice at 9 months after injection.

Survival was improved significantly in both adult- and neonatally-injected groups. Sustained growth improvement was observed in adult-injected mice (Fig. 3C) with a peak of growth velocity in the 2 weeks following the injection of gene therapy (Supplementary Fig. 4B). In neonatally-injected mice, a significant improvement of growth was transiently observed until day 30 (Fig. 3D) consistent with a growth speed similar to WT animals until day 15 (Supplementary Fig. 4B). Later in follow-up, no significant difference of weight was observed between the surviving untreated and neonatally-treated *Asl*^*Neo/Neo*^ mice (Fig. 3E).

A specific fur pattern with sparse, brittle hair called *trichorrhexis nodosa* was observed in untreated *Asl*^*Neo/Neo*^ mice, mimicking symptoms observed in ASA patients ^26^. In adult-injected mice, growth and fur pattern dramatically and sustainably improved compared to untreated *Asl*^*Neo/Neo*^ mice (Figures 3F-H). The correction of the fur phenotype was observed within 2 weeks of gene therapy (Supplementary Fig. 5A); the hair shaft had an improved straighter, more regular shape, a wider medulla and the restoration of the ability to grow and form physiological tips (Supplementary Fig. 5B-D). Fur aspect and growth were improved transiently in neonatally-treated *Asl*^*Neo/Neo*^ mice in the first month of life (Fig. 3I-L).

**Figure 3.**
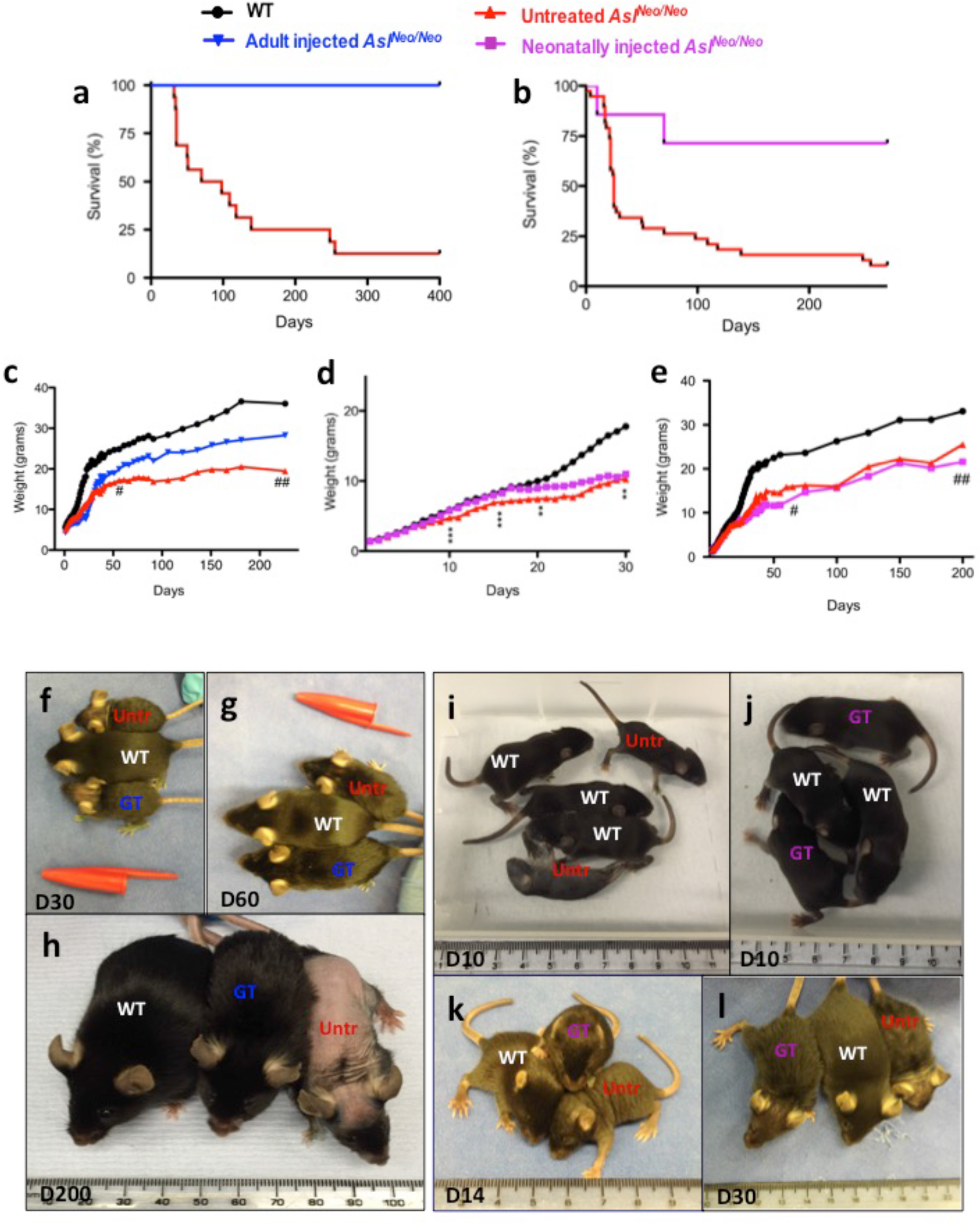
Systemic injection of *AAV8.EFS.mAsl* sustainably improves the macroscopic phenotype of *Asl*^*Neo/Neo*^ mice injected as adults but only transiently after neonatal injection. **(a)** Survival curve of adult-injected *Asl*^*Neo/Neo*^ mice (n=5/5) compared to untreated *Asl*^*Neo/Neo*^ mice (n=2/16); Log rank test p=0.003. (b) Survival curve of neonatally injected *Asl*^*Neo/Neo*^ mice (n=5/7) compared to untreated *Asl*^*Neo/Neo*^ mice (n=2/21); Log rank test p=0.006. **(c)** Mean growth of adult-injected *Asl*^*Neo/Neo*^ mice (n=5) compared to WT (n=11) and untreated *Asl*^*Neo/Neo*^ mice over 12 months (n=19). **(d, e)** Mean growth of neonatally injected *Asl*^*Neo/Neo*^ mice compared to WT (n=31) and untreated *Asl*^*Neo/Neo*^ mice (n=41) **(D)** during the first month (n=13) and **(e)** over 9 months (n=7). **(f-h)** Images of WT, untreated *Asl*^*Neo/Neo*^ (Untr) and adult-injected *Asl*^*Neo/Neo*^ mice with gene therapy (GT). **(i-l)** Images of WT, untreated *Asl*^*Neo/Neo*^ (Untr) and neonatally injected *Asl*^*Neo/Neo*^ mice (GT). Unpaired 2-tailed Student’s *t* test ** p<0.01, *** p<0.001, neonatally injected vs untreated *Asl*^*Neo/Neo*^ mice. # 30% and ## <15% of untreated *Asl*^*Neo/Neo*^ mice still alive; scale bar **(f-l)**: 1 cm.

### Robust long-term correction of the urea cycle in adult- but not neonatally-treated Asl^Neo/Neo^ mice

At 2 months of age, plasma ammonia was similar to that of WT in both adult- and neonatally-injected mice (Fig. 4A). Normal ammonia values persisted until sacrifice at 12 months and 9 months after injection in adult- and neonatally-injected mice, respectively (Fig. 4B). The plasma concentration of argininosuccinic acid was significantly decreased in adult- but not neonatally-injected mice (Fig. 4C). These results were sustained overtime until harvest (Fig. 4D). Similarly, arginine and citrulline plasma concentrations were normalised in adult-injected but not neonatally-injected mice (Supplementary Fig. 6A, 6B). Liver ASL activity was 14.5% of WT activity in untreated *Asl*^*Neo/Neo*^ mice. This increased significantly to 47% and 18.5% in adult- and neonatally-injected groups, respectively, at the time of harvest (Fig. 4E). Anti-ASL liver immunohistochemistry showed a diffuse transduction of cells morphologically identified as hepatocytes, prominent following adult injection and scarce after neonatal injection (Fig. 4F). Quantification of anti-ASL immunohistochemistry showed a significant increase in adult-injected mice (Fig. 4G). Quantitative PCR confirmed greater persistence of vector genomes after adult versus neonatal gene therapy (Supplementary Fig. 6C).

**Figure 4.**
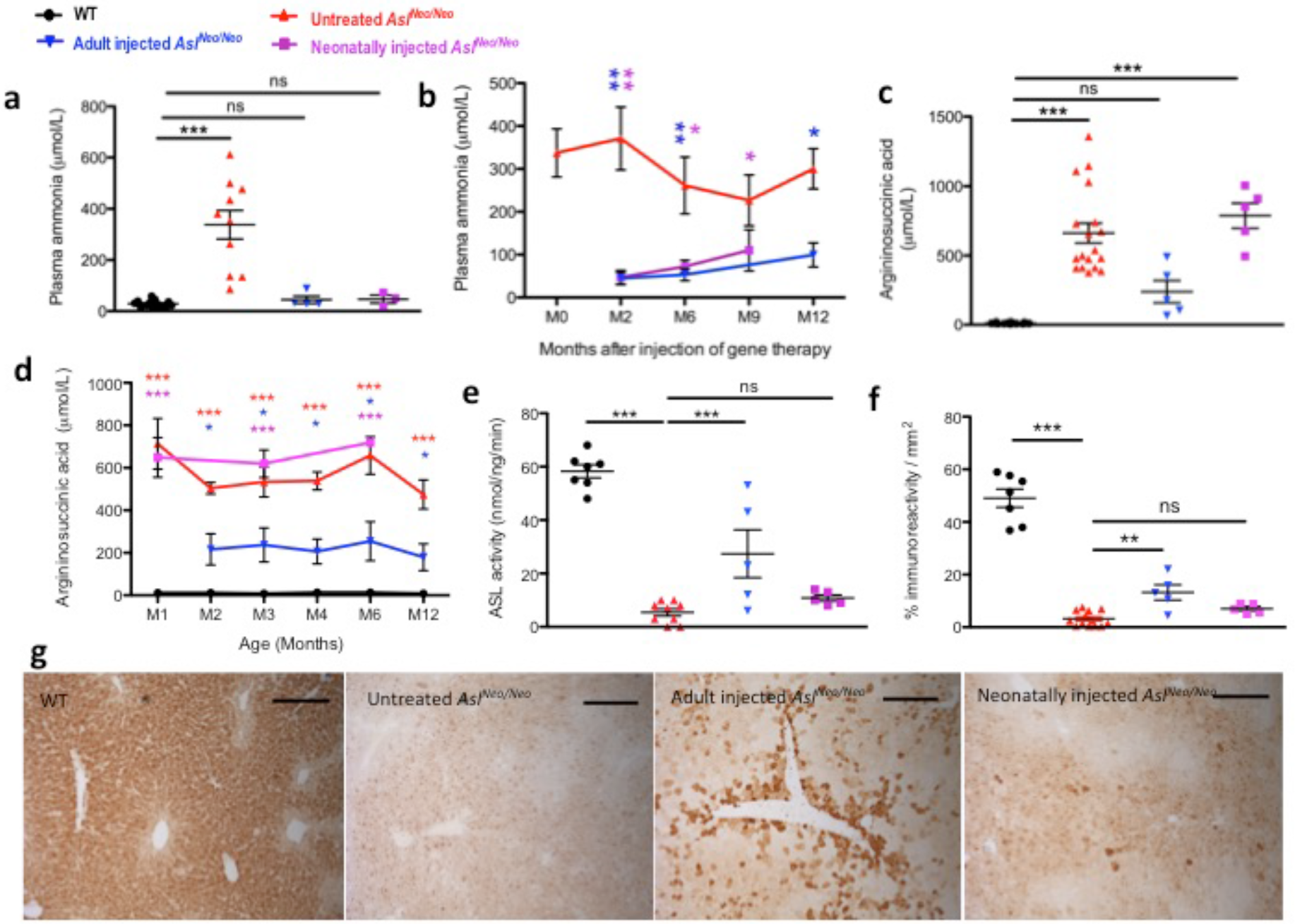
AAV8.EFS.*mAsl* controls hyperammonaemia and restores a functional urea cycle. **(a)** Concentration of plasma ammonia in 2 month-old mice and **(b)** evolution overtime. **(c)** Concentration of argininosuccinic acid in dried blood spots in 2 month-old mice and **(d)** evolution of overtime. **(e)** Liver argininosuccinate lyase (ASL) activity. **(f)** Computational quantification of ASL immunostaining. **(g)** Representative images of ASL-stained sections of liver in WT, untreated, adult-injected and neonatally-injected *Asl*^*Neo/Neo*^ mice (n=5). Horizontal lines display the mean ± SEM. One-way ANOVA with Dunnett’s post-test compared to WT **(a, b)** and untreated *Asl*^*Neo/Neo*^ **(e)**. ns - not significant; * p<0.05; ** p<0.01; *** p<0.001. Scale bars 500 p,m (g); WT n=13-20; untreated *Asl*^*Neo/Neo*^ n=10-12; adult-injected *Asl*^*Neo/Neo*^ n=4-5; neonatally-injected *Asl*^*Neo/Neo*^ n=3-5.

Urinary orotic acid levels were increased significantly in *Asl*^*Neo/Neo*^ mice at 10 weeks of age compared with WT mice. Orotic acid concentration was normalised in two adult-injected mice at 10 weeks however it did not reach statistical significance neither in the adult- nor the neonatally-treated groups (Supplementary Fig. 6D). Plasma alanine amino transferase (ALT) levels were normalised in both adult- and neonatally-injected mice (Supplementary Fig. 6E). Haematoxylin and eosin (H&E) staining of liver samples showed vacuolated cytoplasm in untreated *Asl*^*Neo/Neo*^ mice; cytoplasmic glycogen deposits were identified by periodic acid Schiff (PAS) staining. This feature was markedly improved following adult, but not neonatal injections (Supplementary Fig. 7).

### Long-term improvement of the NO metabolism in the liver

Liver NO levels assessed by nitrite/nitrate levels were reduced in untreated *Asl*^*Neo/Neo*^ mice. These improved in adult-injected but not neonatally-injected mice (Supplementary Fig. 8A). Reduced liver glutathione was decreased in untreated *Asl*^*Neo/Neo*^ mice (Supplementary Fig. 8B).

### Long-term correction of the cerebral NO metabolism in neonatally- but not adult-treated *Asl*^*Neo/Neo*^ mice

Cortical ASL enzyme activity in untreated *Asl*^*Neo/Neo*^ mice was 14.1% of WT activity. In mice injected as adults, this activity was unchanged (16.2% of WT activity) but increased dramatically in mice injected neonatally with 64.8% of WT activity being evident at time of culling (Fig. 5A).

**Figure 5.**
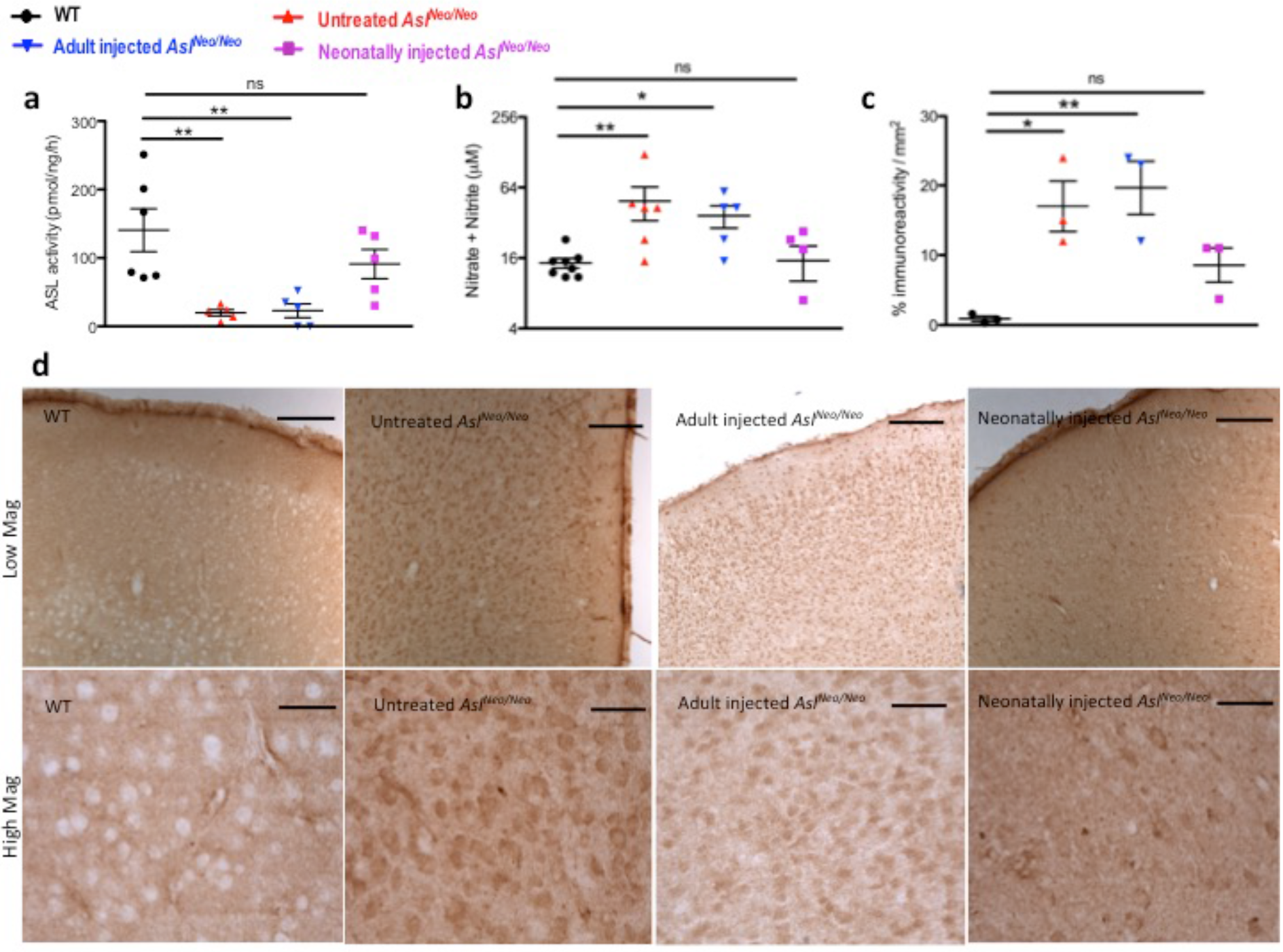
Systemic neonatal but not adult injections of AAV8.EFS.*mAs/* exhibit longterm correction of the impaired NO metabolism in the brain. **(a)** Cortical argininosuccinate lyase (ASL) activity. **(b)** Levels of nitrite/nitrate levels in brain homogenates. **(c)** Computational quantification of nitrotyrosine immunostaining. **(d)** Representative images of nitrotyrosine-stained sections of cortex in WT, untreated, adult-injected and neonatally-injected *Asl*^*Neo/Neo*^ mice (n=3). Horizontal lines display the mean ± SEM. **(a, b, c)** One-way ANOVA with Dunnett’s post-test compared to WT: ns - not significant; 1 p<0.05; ** p<0.01. **(b)** Graph displays not transformed data. Statistical analysis used log-transformed data. WT n=6-8; untreated *Asl*^*Neo/Neo*^ n=5-6; adult-injected *Asl*^*Neo/Neo*^ n=5; neonatally-injected *Asl*^*Neo/Neo*^ n=5. Scale bars of low and high magnification: 500 and 125 μm, respectively.

To assess the effect of the improved ASL activity on the NO metabolism in brains of neonatally-treated *Asl*^*Neo/Neo*^ mice, we measured nitrite/nitrate levels. Compared to WT brains, nitrite/nitrate levels were increased in untreated *Asl*^*Neo/Neo*^ mice and in adult-injected mice by 3.4 and 2.5 times, respectively, whereas in neonatally-injected *Asl*^*Neo/Neo*^ mice the levels were not significantly different from WT mice (Fig. 5B). To examine if this decrease in nitrite/nitrate levels in neonatally-treated mice was correlated with a modification of the oxidative/nitrosative stress, we quantified cortical nitrotyrosine staining. There was no significant difference between neonatally-injected mice and WT mice. In contrast, adult injected mice and untreated *Asl*^*Neo/Neo*^ mice showed a significant increase in the percentage of immunoreactivity (Fig. 5C, 5D).

### Impact of gene therapy on behaviour and cell death in the brain

Behavioural tests were performed to assess open field exploration. At 3 months of age, there was a significant reduction in the walking distance measured in the untreated *Asl*^*Neo/Neo*^ mice, whereas an improvement was seen in both adult- and neonatally-injected groups (Fig. 6A). Performance with an accelerating rotarod at the same age was significantly reduced in untreated *Asl*^*Neo/Neo*^ mice but not significantly different from WT in both adult- and neonatally-injected groups (Fig. 6B).

**Figure 6.**
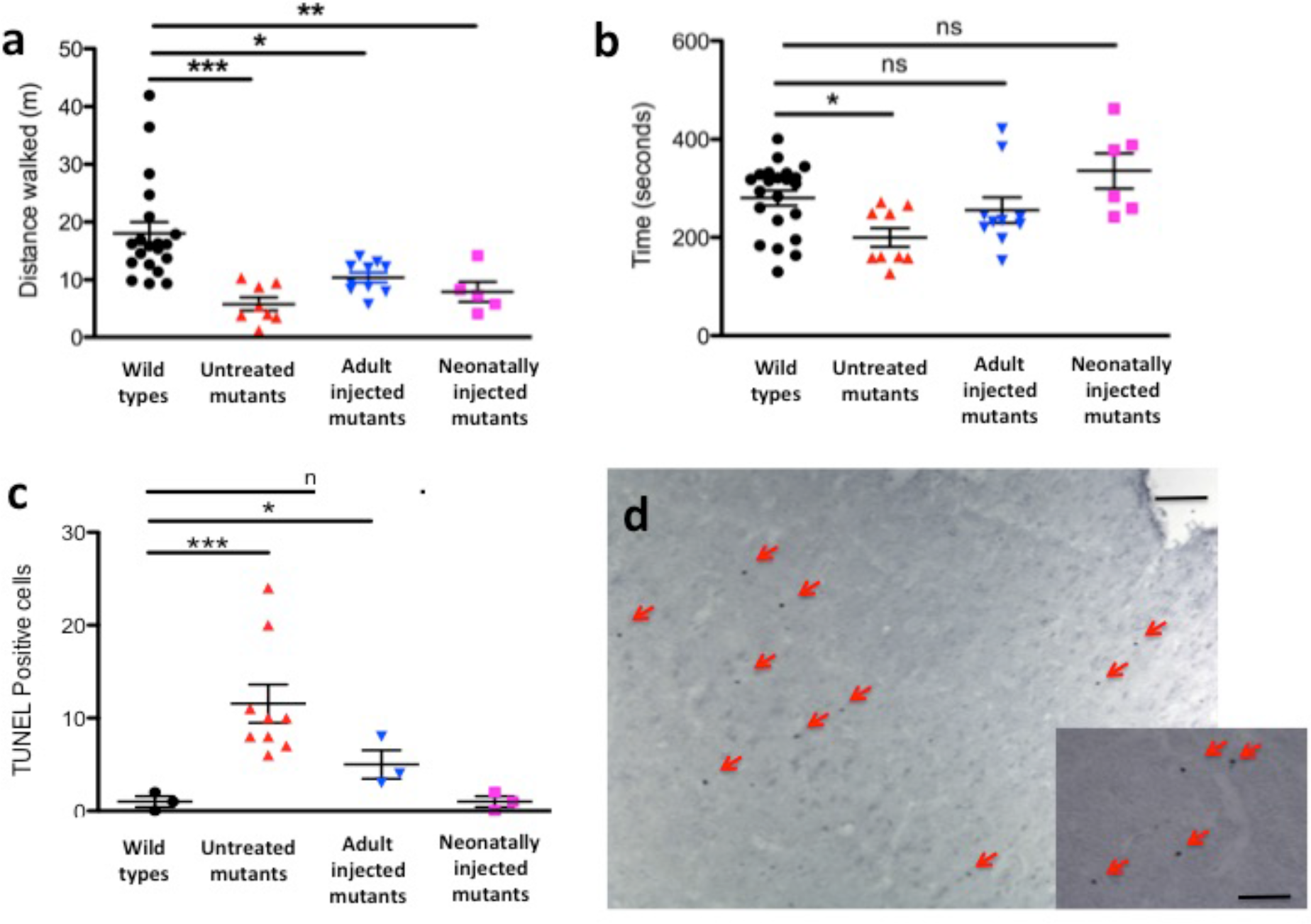
AAV-mediated gene therapy improves behavioural testing and is inversely correlated with cortical neuronal loss in treated animals. **(a)** Open field test and **(b)** accelerating rotarod performed in 2 month-old mice. **(c, d)** Apoptotic cells at time of culling in WT (aged 9 - 12 months), untreated (aged 21 days - 12 months), adult-injected (aged 12 months) and neonatally-injected *Asl*^*Neo/Neo*^ mice (aged 9 months). (c) Cell counting in cortex. **(d)** Representative cortex image of an untreated 25 days old *Asl*^*Neo/Neo*^ mouse symptomatic of hyperammonaemia and culled for humane endpoint. Horizontal lines display the mean ± SEM. **(a, b)** One-way ANOVA with Dunnett’s post test compared to WT: ns - not significant, *p<0.05, ** p<0.01, *** p<0.001. **(c)** Graph displays not transformed data. Statistical analysis used log-transformed data. WT n=20-22; untreated *Asl*_*Neo/Neo*_ n=8-9; adult-injected *Asl*_*Neo/Neo*_ n=5; neonatally-injected *Asl*_*Neo/Neo*_ n=5. Scale bars: 500 μm and 125 μm in main and inset pictures, respectively.

Cell death was assessed by TUNEL staining and was found to be significantly increased in the cortex of untreated *Asl*^*Neo/Neo*^ mice compared to WT. Cell death was reduced in adult-injected and neonatally-injected mice compared to untreated *Asl*^*Neo/Neo*^ mice (Fig. 6C, 6D).

### DISCUSSION

This study provides new insight into the pathophysiology of the brain disease in ASA and presents proof-of-concept of hepatocerebral phenotypic correction of the hypomorph *Asl*^*Neo/Neo*^ mouse model *via* systemic AAV-mediated gene therapy.

Compared to other urea cycle defects, the neurological disease in ASA is a paradox between low rates of hyperammonaemic decompensation and high rates of neurological complications, including neurocognitive delay, abnormal neuroimaging, epilepsy and ataxia ^18^. The *Asl*^*Neo/Neo*^ mouse model was used to study its unreported neurological phenotype and investigate neuropathophysiology. NO plays a complex and ambiguous role in the brain, involved in both inflammation-related neurotoxicity and cGMP-mediated neuroprotection ^27,28^. Increased formation of NO (assessed by nitrite/nitrate levels) and cGMP concentrations in the brain of *Asl*^*Neo/Neo*^ mice suggests a persisting physiological up-regulation of the NO/cGMP pathway, which is observed with appropriate coupling of NOS ^20^. However previous studies in *Asl*^*Neo/Neo*^ mice have demonstrated the uncoupling of NOS likely promoted by low tissue L-arginine content ^29^, which is associated with the increase of systemic biomarkers of oxidative stress ^21^. NOS uncoupling causes oxidative/nitrosative stress *in vitro* with the production of peroxynitrite, which in turn contributes to decrease of antioxidants, inhibition of the mitochondrial respiratory chain, opening of the permeability transition pore and cell death ^24^. This is consistent with our observation of increase of both neuronal nitrotyrosine staining and nitrite/nitrate levels. In the brain, hyperammonaemia increases nNOS and iNOS-mediated NO synthesis *via* an increase in extracellular glutamate and oxidative stress ^25^. In our study, correction of hyperammonaemia only did not modify the oxidative/nitrosative stress, suggestive of an independent brain-specific causative mechanism. A measurable benefit in cell death in the cortex is observed when neuronal ASL activity is restored. Neurons are more vulnerable to oxidative stress than astrocytes *in vitro*, as they cannot overexpress γ-glutamyl transpeptidase to replenish their intracellular glutathione content ^30^. Therefore, they rely on the paracrine glutathione supply from astrocytes when exposed to reactive nitrogen species and oxidative stress ^30^. This might explain the neuronal staining observed for nitrotyrosine and the efficacy of neuronal-targeted gene therapy. This nitrosative stress caused by peroxynitrite, generating nitrotyrosine, has been implicated previously in the pathophysiology of various neurodegenerative diseases: Parkinson’s disease ^31^, Alzheimer’s disease ^32,33^ and amyotrophic lateral sclerosis ^34^. In urea cycle defects, a neuronal disease caused by oxidative stress as a consequence of low tissue arginine has been hypothesised as playing a role in the brain pathophysiology ^35^. Although the precise biochemical mechanisms regulating NO metabolism in different cerebral cell types in ASA remain elusive, NOS coupling and uncoupling appear to co-exist in the brain accounting for both the physiological NO-cGMP pathway and nitrosative/oxidative stress, respectively (Fig. 7) as observed in Alzheimer’s disease ^27^. No evidence of neuroinflammation was observed in astrocytes and microglial cells, when GFAP and CD68 were assessed by immunohistochemistry suggesting that these cell types are not primarily involved. Thus neuronal oxidative/nitrosative stress seems to play a key-role in the ASA brain disease.

**Figure 7.**
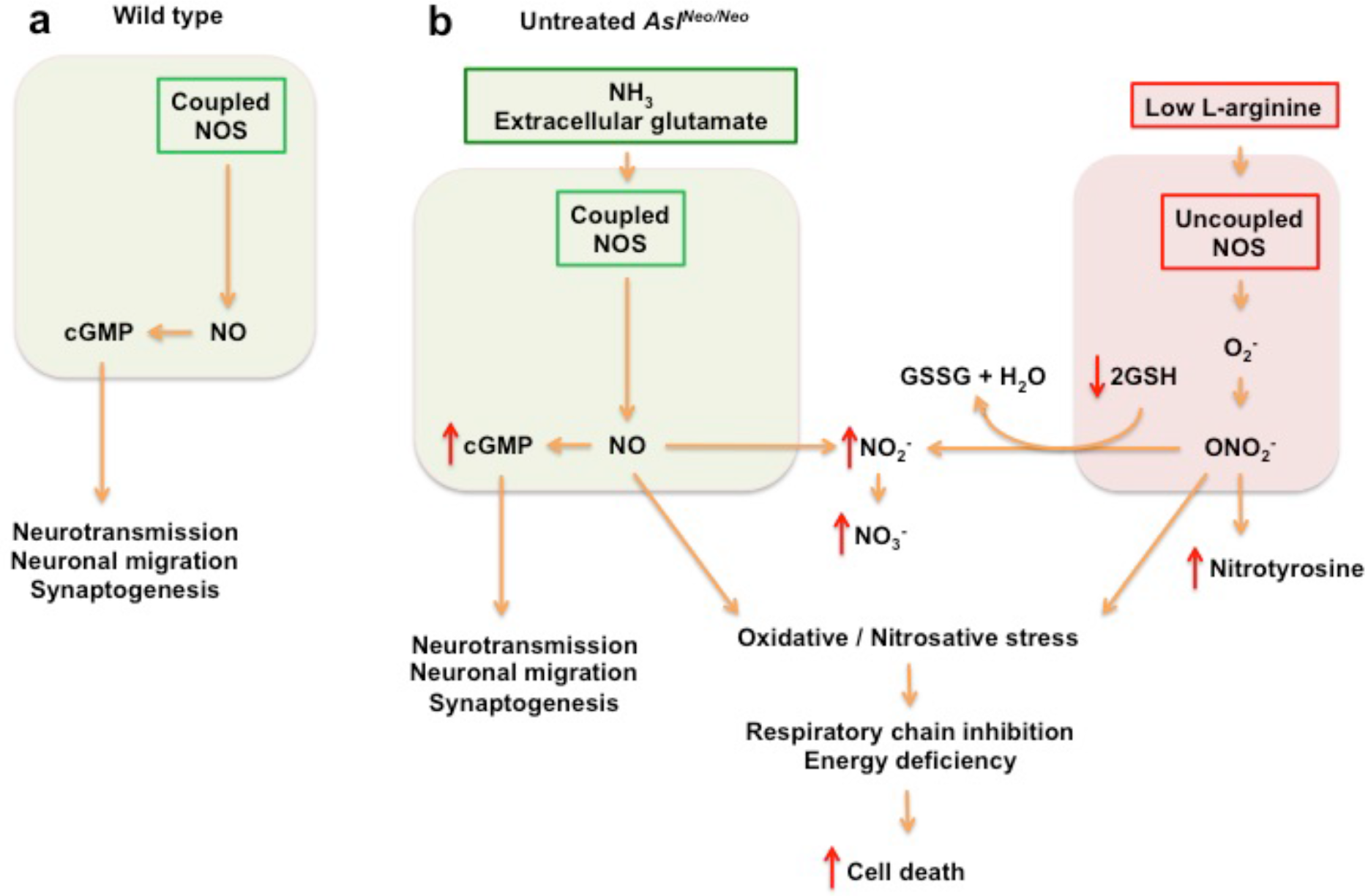
Proposed pathophysiology of the brain disease in *As/^Neo/Neo^* mice. **(a)** Residual coupled NOS produces NO with increased cGMP production. **(b)** Low L-arginine causes NOS uncoupling and produces superoxide ions (O_2_^−^), which will generate the production of peroxynitrite (ONOO^−^). ONOO^−^ causes oxidative/nitrosative stress with production of nitrotyrosine and nitrite (NO_2_^−^) after detoxification of ONOO^−^ by reduced glutathione (GSH). Oxidative/nitrosative stress will impair the respiratory chain, alter the energy production of the cell and lead to cell death. Neurons are particularly vulnerable as they cannot increase their production of glutathione adequately, and rely on an astrocytic supply. **(b)** Red arrows represent modified measured parameters in *Asl*^*Neo/Neo*^ compared to WT mice. cGMP: cyclic guanosine monophosphate, GSH: reduced glutathione, GSSG: oxidised glutathione, NO: nitric oxide, NO_2_^-^: superoxide ion, ONO_2_^−^: peroxynitrite, NOS: nitric oxide synthase.

In murine animal models, AAV8 is known for its ability to widely transduce the brain after intracranial administration ^36,37^. After systemic injection, most of the organs and especially the liver are successfully targeted ^38^ but the neurotropism is influenced by the age of infusion and the dose of vector administered. For instance, successful brain transduction with AAV8 and a CMV promoter after systemic delivery has been reported previously until day 14 of life but not later (1.5×10^11^ vg/mouse) ^39^. However brain transduction was barely detectable in adult mice after a similar experiment (1×10^11^ vg/mouse) ^38^. Increasing the dose of vector improved brain transduction in adult mice. Indeed intravenous injection in adult mice with AAV8 and EF1α promoter showed mild brain cell transduction at 1×10^11^ to 2×10^12^ vg/mouse ^40,41^, but widespread neuronal and astrocytic transduction at 7.2×10^12^ vg/mouse (approximately 2.9×10^14^ vg/kg) ^42^. The transient ability for AAV vectors to cross an immature blood brain barrier in the neonatal mouse brain is not well understood and could be due to immaturity or receptor-mediated transcytosis ^39,43^. The age at injection does not only allow an increased brain transduction but influences the cell types transduced. A predominant neuronal transduction is observed when the vector is administered during the first 48 hours of life whereas a preferential astrocytic transduction is noticed from day 3 onwards ^36^. These observations were made with an AAV9 serotype, in mice ^44,45^ and non-human primates ^46,47^, with a long-standing transgene expression of up to 18 months in mice after a single systemic neonatal injection ^39^. For this study, the choice of EFS promoter was made on the basis of various advantages: relatively ubiquitous expression and strong promoter activity ^48^, resistance to silencing ^49^, reduced potential risk of insertional mutagenesis compared to other ubiquitous promoters ^50^, already used in clinical trials ^51^. In the brain, the AAV8.EFS. *GFP* vector targeted mainly neurons due to its promoter specificity ^52^ and the neonatal timing of injection ^36^.

Long-term correction of ammonaemia levels was observed in mice injected as adults or neonates demonstrating successful restoration of ureagenesis. Adult-injected mice exhibited a more complete correction with prolonged improvement of other typical features of the deficient urea cycle: growth, fur, blood amino acid concentrations and in some, orotic aciduria and liver intracellular glycogen deposits. The correction of the phenotype correlated with the liver ASL activity. AAV vectors deliver non-integrating transgene copies, persisting as episomes in the transduced cell. The transient nature of the majority of the metabolic effects after neonatal injection is likely caused by a loss of transgene vector genomes in the rapidly growing liver during the first weeks of life and is consistent with previous studies ^53,54^. These results are in line with previous experiments using AAV-derived vectors in murine models of urea cycle defects ^10,11,13^. In this study, an increase from 14.5% to 18% of WT liver ASL activity was observed 9 months after neonatal injection. This provided a persistent correction of ammonaemia but was not sufficient to normalise other biochemical parameters of the disease (plasma amino acids, orotic aciduria). In ASA, the increased urine secretion of the argininosuccinic acid that removes two nitrogen moieties may explain reduced tendency to develop hyperammonaemic episodes compared to proximal urea cycle defects ^16^. AAV-mediated correction of other models of urea cycle disorders has shown that a small (approximately 3%) improvement in liver enzyme levels and ureagenesis can restore survival and improve ammonia levels ^55,56^. Controlling orotic aciduria in the *Spf*^*ash*^ mouse model of ornithine transcarbamylase deficiency however required 5 times more vector compared to that necessary to normalise ammonaemia ^9,57^. In that respect, our study provides a hierarchization in the biomarkers significance according to the ASL residual activity in ASA. Plasma amino acids and urine orotic acid required a liver ASL activity of >18% for normalization whereas ammonaemia was seen to normalize when ASL activity was only 14.5-18% of WT activity. However these figures might be biased by the persistence of the non-integrating transgene delivered by AAV vector. As reported previously in shRNA-induced hyperammonaemic *Spf*^*ash*^ mice ^9^, the AAV-encoded enzymatic activity required to normalize ammonaemia might be higher than the endogenous residual activity required in a non-hyperammonaemic subject. Indeed the reduced transgenic expression from non-integrated episomes compared to endogenous chromosomal alleles has been suspected recently from results of a liver-directed clinical trial ^58^. The fur phenotype observed in ASL- and argininosuccinate synthase-deficient mice is likely to be caused by hypoargininaemia as arginine represents up to 10% of hair composition ^17^ The long-term phenotypic improvement of the fur in adult-injected mice is consistent with the improved plasma arginine levels.

A small but significant increase of the nitrite/nitrate levels was observed in the livers of adult-injected mice thereby suggesting that restoration of ASL activity had a positive effect on the function of both urea and citrulline-NO cycles. A decrease in reduced glutathione in untreated *Asl*^*Neo/Neo*^ mice reflects increased cellular levels of oxidative stress. Whilst systemic NO deficiency ^15^ and increased oxidative stress ^21^ have been described previously, it was shown that long-term correction of neuronal ASL achieved a marked decrease of the cortical oxidative/nitrosative stress, independent of improvement in ammonia levels.

ASA patients are at high risk of developing neurological complications even if hyperammonaemic episodes do not occur ^18^. Our data provide further evidence for a neuronal disease with oxidative/nitrosative stress independent of ammonaemia, and illustrates the pathophysiological importance of disturbed NO metabolism in the ASA brain. Any therapy aiming to preserve the neurological status of ASA patients needs to protect the brain from two potential insults, hyperammonaemia and disturbed cerebral NO metabolism. Similar to liver transplantation ^59^, a gene therapy approach targeting only hepatocytes will cure the urea cycle defect but will not correct the symptoms related to ASL deficiency in extra-hepatic tissues, especially the brain ^21^. This study provides proof-of-concept for phenotypic correction of the *Asl*^*Neo/Neo*^ mouse model using AAV technology. These promising results raise the possibility of combining two sequential systemic injections: i) a first early (neonatal) injection of a gene therapy vector that would transiently restore the urea cycle in the liver and will transduce neurons to modify the long-term natural course of the neuronal disease, and ii) a second injection in infancy or adulthood targeting the liver for long-term correction of the urea cycle. The potential for humoral immune response generated by the first AAV injection will need to be considered for the second injection ^60^. It is possible however, similarly to what is reported for neonatal rodents ^61^, that the immaturity of the immune system in humans at the time of neonatal injection might prevent the induction or diminish the magnitude of humoral response against the AAV capsid ^62^. An alternative AAV capsid, which does not cross-react with neutralising antibodies developed, might be a valid option ^63,64^ Several inherited metabolic diseases with hepatocerebral phenotype might benefit from a similar dual targeting approach such as mitochondrial diseases caused by nuclear genetic defects (e.g. *POLG1*, *MPV17*, *DGUOK* genes) and some lysosomal storage disorders (e.g. neuronopathic Gaucher disease, mucopolysaccharidosis type I and II). Depending on the pathophysiology of the disease, specific brain cell-types can be selectively targeted in modifying either promoter and/or age at injection ^36,37^.

## METHODS

### Animals

The *Asl*^*Neo/Neo*^ mice (B6.29S7-*Asl*^*tm1Brle*^/J) were purchased from Jackson Laboratory (Bar Harbor, ME). WT or heterozygote mice were maintained on standard rodent chow (Harlan 2018, Teklab Diets, Madison, WI; protein content 18%) with free access to water. *Asl*^*Neo/Neo*^ mice were started on a supportive treatment including a reduced-protein diet (5CR4, Labdiet, St Louis, MO; protein content 14.1%) from day 15 to day 50 and received daily intraperitoneal injections of sodium benzoate (0.1g/kg/d) and L-arginine (1g/kg/d) from day 10 to day 30. C57BL/6 mice were purchased from Jackson Laboratory (Bar Harbor, ME).

### Behavioural testing. Rotarod

After a period of acclimatisation including 3 trials/day for 3 consecutive days, the test was performed on a Rotarod LE 8200 (Panlab, Harvad apparatus, Cambridge, UK) with 3 attempts/day for 5 consecutive days. The latency to fall from the rod under continuous acceleration from 4 to 40 rpm over 5 minutes was recorded. *Open field test*. The animal was placed in the centre of a plastic box (40 × 40 cm floor) and video-recorded for 5 minutes. Computational analysis of the distance walked was performed automatically using the MouseLabTracker v0.2.9 application on Matlab software (Mathworks, Natick, MA, USA) ^65^.

### Genotyping

DNA extraction from tail or ear clips was performed by adding 25 mM NaOH, 0.2 mM EDTA adjusted to pH 12 prior to heating the sample at 95°C for 10 minutes. An equal volume of 40 mM Tris (adjusted to pH 5) was then added. DNA was amplified using a Taq DNA Polymerase PCR kit (Peqlab, Germany) according to manufacturer’s instructions using the following primers: 5’-GGTTCTTGGTGCTCATGGAT-3’ (sense), 5’-GCCAGAGGCCACTTGTGTAG-3’ (WT, antisense) and 5’-CATGACAGCTCCCATGAAGA-3’ *(Asl^Neo/Neo^* mice, antisense) provided by Jackson Laboratory (Bar Harbor, ME). Amplification conditions were 95°C for 10 minutes then 40 cycles of 94°C for 30 seconds, 63°C for 30 seconds, 72°C for 1 minute.

### Cell culture

Human embryonic kidney (HEK) 293 cells were maintained in Dulbecco’s modified Eagle medium (Gibco, Invitrogen, Gran Island, NY) supplemented with 10% (vol/vol) fetal bovine serum (JRH, Biosciences, Lenexa, KS) and maintained at 37°C in a humidified 5% CO2-air atmosphere.

### Virus production and purification

The murine *Asl (mAsl)* gene (Vega Sanger Asl-0003 transcript OTTMUST00000085369) inserted in a pCMV-SPORT6 gateway vector was purchased from Thermo Scientific (Loughborough, UK). A single stranded AAV2 backbone plasmid containing an expression cassette with the elongation factor 1 α short promoter (EFS), a modified simian virus 40 (SV40) small t antigen intron, the human vacuolar protein sorting 33 homolog *(hVPS33B)* cDNA, Woodchuck hepatitis post regulatory element (WPRE) sequence, SV40 late polyA (courtesy of P. Gissen) was digested with EcoRV-Nhe1 to remove the *hVPS33B* cDNA. Subsequently the *mAsl* cDNA, was digested with EcoRV-Nhe1 and ligated into this vector. Vector production was performed by triple transfection in HEK293T cells following polyethylenimine transduction as described previously ^66^. Vector purification was performed by affinity chromatography on an ÄKTAprime plus (GE Healthcare UK Ltd, Buckinghamshire, UK) with Primeview 5.0 software with a HiTrap AVB Sepharose column (GE Healthcare UK Ltd, Buckinghamshire, UK) according to manufacturer’s instructions. Vector quantification was performed by electrophoresis on an alkaline gel ^67^. An AAV8 vector encapsidating a single stranded DNA sequence containing the *GFP* gene under the transcription activity of the EFS promoter, the SV40 intron upstream and WPRE and polyA downstream *GFP* was provided by J. Hanley.

### Stereoscopic fluorescence microscopy

At 5 weeks of age, C57BL/6 mice injected intravenously at day 0 with 1.7×10^11^vg/mouse of AAV8.EFS.*GFP* and control littermates were culled by terminal exsanguination and perfused with PBS. GFP expression was assessed using a stereoscopic fluorescence microscope (MZ16F; Leica, Wetzlar, Germany). Representative images were captured with a microscope camera (DFC420; Leica Microsystems, Milton Keynes, UK) and software (Image Analysis; Leica Microsystems).

### Free-floating and paraffin embedded immunohistochemistry

Brain tissue was fixed in 4% paraformaldehyde (PFA) over 48 hours then stored in 30% sucrose at 4°C. Cryo-sectioning was performed with a Microm freezing microtome (Carl Zeiss, Welwyn Garden City, UK). Immunohistochemistry was performed on free-floating sections blocked in Tris-buffered saline-Triton (TBST)/15% normal goat serum and incubated overnight at 4°C using anti-GFP (1:10,000; Ab290, Abcam, Cambridge, UK), anti-nitrotyrosine (1:800; 06-284, Merck Millipore, Temecula, CA, USA), anti-nNOS (1:200; bs10197R, Bioss antibodies, Woburn, MA, USA), anti-iNOS (1:500; NBP1-50606, Novus Biologicals, Abingdon, UK), anti-eNOS (1:300; 610296, BD Transduction Lab), anti-GFAP (1:500; MAB3402, Merck Millipore, Temecula, CA, USA), anti-CD68 (1:100; MCA1957, Biorad, Serotec) antibodies diluted in TBST/10% normal goat serum. After 3 washes with Tris-buffered saline (TBS), a 2 hour incubation with a biotinylated goat anti rabbit secondary antibody (1:1000; Vector, Burlingame, CA, USA) at room temperature was followed by 3 TBS washes before a 2 hour incubation with avidin-biotinylated horse radish peroxidase (ABC; 1:100; Vector, Peterborough, UK) at room temperature. After 3 TBS washes, detection was performed with a 0.05% 3,3’-diaminobenzidine (DAB) solution diluted in TBS. The reaction was stopped by the addition of ice cold TBS. Three TBS washes were performed before mounting the tissue on chrome-gelatin-coated slides. Slides were cover-slipped with DPX-new (Merck Millipore Corporation, Temecula, CA, USA).

Systemic organs were fixed with 10% formalin for 48 hours and stored in 70% ethanol at 4°C. Paraffin embedded sections were dewaxed, dehydrated in an ethanol gradient. Blocking was performed with 1% hydrogen peroxide in methanol for 30 minutes followed by antigen retrieval using 10 mmol/l sodium citrate buffer pH 7.4. Non-specific binding was blocked with 15% normal goat serum (Vector, Burlingame, CA, USA). After 3 washes in PBS, sections were incubated with anti-ASL (1:1000; Ab97370, Abcam, Cambridge, UK) and anti-GFP (1:1000; Ab290, Abcam, Cambridge, UK) antibodies overnight at 4 °C. Detection was performed with Polink-2 HRP Plus Rabbit Detection System for Immunohistochemistry (GBI labs, Mukilteo, WA, USA) as per manufacturer’s instructions. After dehydration in a gradient of ethanol and three washes in xylene, slices were cover-slipped with DPX-new (Merck Millipore Corporation, Temecula, CA, USA). Images were captured with a microscope camera (DFC420; Leica Microsystems, Milton Keynes, UK) and software (Image Analysis; Leica Microsystems).

### Free-floating immunofluorescence

Free-floating sections were blocked in TBST/15% normal goat serum and incubated overnight at 4°C with anti-GFP (1:4000; Ab290, Abcam, Cambridge, UK) and anti-NeuN (1:4000; Millipore, Billerica, MA, USA) diluted in TBST/10% normal goat serum as described previously ^44^ After 3 washes with TBS, samples were incubated for 2 hours with goat anti-rabbit Alexa 488 and goat anti-mouse Alexa 546 (1:1000; Invitrogen, Paisley, UK). After a further 3 washes, sections were incubated with DAPI (1:2000; Invitrogen) and mounted on chrome-gelatin-coated slides and cover-slipped with Fluoromount (Southern Biotech, Birmingham, AL, USA).

### TUNEL staining

TUNEL staining was performed as described previously ^68^ using the Roche kit (Roche, Welwyn Garden City, Hertfordshire, UK). Briefly, sections were incubated in 3% hydrogen peroxide in methanol for 15 min and washed in 0.1 M phosphate buffer (PB) before incubation with terminal deoxytransferase (TdT) and deoxyuridine trisphosphate (dUTP) in a solution of 0.1% TdT, 0.15% dUTP, 1% cacodylate buffer) at 37°C for 2 hours. The reaction was stopped by incubating the section in TUNEL stop solution (300 mM NaCl, 300 mM sodium citrate) for 10 min. Sections were then washed in 3 × 0.1 M PB solution, incubated with avidin-biotinylated horse radish peroxidase (ABC; 1:100; Vector, Peterborough, UK) at room temperature for 1 hour, washed 4 times in 10 mM PB and visualised with DAB enhanced with cobalt nickel. The reaction was stopped in 10 mM PB and washed twice in double-distilled (ddH2O) water.

### Nissl staining

Brain sections were fixed in 4% PFA for 24 hours then in 70% ethanol for 24 hours. On day 3, sections were incubated in Cresyl Violet solution (BDH, East Grinstead, West Sussex, UK) for 3 minutes followed by dehydration in an ethanol gradient (70%, 90%, 96%, 96% with acetic acid, 100%), isopropanol, and three washes in xylene before being cover-slipped with DPX-new (Merck Millipore Corporation, Temecula, CA, USA).

### Quantitative analysis of immunological staining

Ten random images per sample were captured with a microscope camera (DFC420; Leica Microsystems, Milton Keynes, UK) and software (Image Analysis; Leica Microsystems). Quantitative analysis was performed with thresholding analysis using the Image-Pro Premier 9.1 software (Rockville, MD, USA).

### Blood chemistry

Plasma ammonia and alanine aminotransferase (ALAT) were analysed by Chemical Pathology Great Ormond Street Hospital, London

### Mass spectrometry

Blood was spotted onto a Guthrie card and allowed to dry at room temperature for 24 hours. Amino acids were measured in dried blood spots by liquid chromatography-tandem mass spectrometry (LC-MS/MS). A 3mm-diameter punch was incubated for 15 min in a sonicating water bath in 100 μL of methanol containing stable isotopes used as internal standards (2 nmol/l of L-Arginine-13C (CK isotopes, Ibstock, UK) and L-Citrulline-d7 (CDN istotopes, Pointe-Claire, Quebec, Canada). A 4:1 volume of methanol was added to precipitate contaminating proteins. The supernatant was collected and centrifuged at 16,000 g for 5 minutes. Amino acids were separated on a Waters Alliance 2795 LC system (Waters, Midford, USA) using a XTerra^®^ RP18, 5 μm, 3.9 × 150mm column (Waters, Midford, USA). The mobile phases were (A) methanol (B) 3.7% acetic acid. The gradient profile using a constant flow rate of 0.2mL/min, with initially 100% B for the first minute and gradually increasing the flow of A as follows: 85% from 1 to 6 minute, 75% from 6 to 8 minute, 5% from 9 to 15 minute, 100% from 16 to 25 minute. The column was reconditioned for 10 minutes at the end of each run. Detection was performed using a tandem mass spectrometer Micro Quattro instrument (Micromass UK Ltd, Cheshire, UK) using multiple reaction monitoring in positive ion mode and ion transitions published previously (279). The temperature of the source and for desolvation were 120°C and 350°C, respectively. The capillary and cone voltages were 3.7 kV and 35 V, respectively. The cone gas flow was 50 L/h and the syringe pump flow 30 μL/min. The mass spectrometer vacuum was 4.3×10^−3^ mbar. The multiplier and extractor voltages were 650 V and 1 V, respectively. Data were analysed using Masslynx 4.1 software (Micromass UK Ltd, Cheshire, UK). Calibration curves ranging between 0 to 500 μM were constructed to enable quantification.

### Analysis of ASL enzyme activity

Liver and brain samples were snap-frozen in dry ice at time of collection after perfusion of the animal with PBS to remove residual blood in tissues. Protein extraction was performed on ice. Samples were homogenised in lysis buffer (50 mM Tris, 150 mM NaCl, 1% Triton adjusted to pH 7.5) and centrifuged at 16,000 g for 20 min at 4°C. Protein quantification of the supernatant was performed using the Pierce™ BCA protein assay kit (ThermoFisher Scientific, Rockford, IL, USA) according to manufacturer’s instructions.

Liver ASL activity was measured in duplicate samples. 20μg protein was added to a buffer solution of 20 mM Tris, 1 mM argininosuccinic acid, 0.02 nM L-citrulline-d7 and incubated for 2 hours at 37°C. Brain ASL activity was also measured in duplicate samples. 80μg protein was added to a buffer solution of 20 mM Tris, 30 μM argininosuccinic acid, 0.02 nM L-citrulline-d7 and incubated for 2 hours at 37°C. A 4:1 volume of methanol was added to stop the reaction, and centrifuged at 9,500 g for 2 minutes. The supernatant was analysed by the LC-MS/MS method described above. ASL activity was calculated by subtracting the amount of argininosuccinic acid post-incubation from that pre-incubation.

### Nitrite and nitrate measurement

Measurement of nitrite and nitrate levels were performed using a modified Griess reaction protocol ^69^. Samples were collected carefully in order to minimise the risk of nitrite and nitrate contamination. All glassware and plastic ware were cleaned with double-distilled water (ddH2O). Animals were anaesthetised and perfused on ice. Organs and brain were collected on ice in less than 3 minutes and snap frozen on dry ice. Samples were homogenised in 2 volumes of ddH_2_O with a grinder (Tissue Master 125, OMNI International, Kennesaw, GA, USA) on ice and then centrifuged at 13,500 g at room temperature for 10 minutes in 3,000 kDa cut-off filters (Merck Millipore, Darmstadt, Germany). Enzyme stocks were made and stored as follows: Nitrate reductase was dissolved in 1:1 phosphate buffer:glycerol to a final specific activity of 59 units/mL and stored at −20°C. Glucose-6-phosphate dehydrogenase was dissolved in 1:1 phosphate buffer:glycerol to a final specific activity of 125 units/mL and stored at −20°C. The enzyme master mix was made fresh for each experiment using a mixture of 1 mL of phosphate buffer, 10 μL of nitrate reductase stock solution, 10 μL of glucose-6-phosphate dehydrogenase stock solution and 1.14 mg of glucose-6-phosphate. A calibration curve, covering a range of 0 - 600 μmol/l was prepared using serial dilutions of nitrite and nitrate standards. Analysis was performed in a 96-well plate and each sample analysed in duplicate. 60 μL of sample was mixed with 10 μL of 10 mmol/l NADPH and 40 μL of enzyme mastermix. Samples were incubated for 1 hour at room temperature on a rotating shaker to allow conversion of NO_3_” to NO_2_”. The Griess reaction was performed by adding 75 μL of 116 mM sulfanilamide, 5% phosphoric acid, then 75 μL of 7.7 mM N-1-naphtylethylenediamine dihydrochloride. The plate was read at 550 nm in a FLUOstar Omega spectrophotometer (BMG Labtech, Ortenberg, Germany).

### cGMP measurement

Brain samples from PBS perfused mice were flash-frozen in dry ice and ground in liquid nitrogen. Samples were weighed and diluted in 10 volumes of 0.1M HCl prior to centrifugation at 600 g for 10 minutes. cGMP was measured using a cGMP complete ELISA kit (Enzo Life Sciences, Farmingdale, NY, USA) according to manufacturer’s instructions in a non-acetylated format reaction. Samples were analysed in a 96-well plate and were analysed at 405 nm in a FLUOstar Omega spectrophotometer (BMG Labtech, Ortenberg, Germany).

### Glutathione analysis

Reduced glutathione was measured as described previously ^70^. Briefly snap-frozen samples were homogenized in 20 volumes of cold 50 mM Tris buffer pH 7.4 and sonicated. Monochlorobimane (at a final concentration of 100 μM) and glutathione-S-transferase (1 U/mL) were added to the samples prior to incubation at room temperature whilst protected from light for 30 minutes. Samples were analysed in a FLUOstar Omega spectrophotometer (BMG Labtech, Ortenberg, Germany) using an excitation wavelength of 360 nm and an emission wavelength of 450 nm.

### Green fluorescent protein ELISA

Lysis buffer was added to cover the sample prior to grinding with a mixer after a couple of freeze/thaw cycles. The sample was then centrifuged for 2 minutes at 1,200 g and the supernatant collected. The protein concentration for each sample was measured using the Pierce™ BCA protein assay kit (ThermoFisher Scientific, Rockford, IL, USA) according to manufacturer’s instructions and read at 570 nm in a FLUOstar Omega spectrophotometer (BMG Labtech, Ortenberg, Germany). Each step of the ELISA protocol was separated by 3 washes of a wash buffer 0.05% Tween20 in PBS. A monoclonal anti-GFP antibody (1:10,000; ab1218, Abcam, Cambridge, UK) was added, the plate sealed, incubated overnight at 4°C then blocked with 1% bovine serum albumin in PBS for 1 hour at 37°C. Analysis was done in a sealed 96-well plate and all samples were measured in duplicate. GFP standards were serially diluted in wash buffer and incubated alongside a buffer blank and the samples for 1 hour at 37°C. An anti-GFP biotin-conjugated secondary antibody (1:5,000; Ab6658, Abcam, Cambridge, UK) followed by a streptavidin-horseradish peroxidase conjugate (1:20,000; SNN2004, Invitrogen, Camarillo, CA, USA) were added to the samples. Both were incubated for 1 hour at 37°C successively. Tetramethylbenzidine (TMB) was then added for 2 minutes at room temperature before the reaction was stopped with 2.5 M H2SO4. The plate was read at 450 nm within 30 minutes of having stopped the reaction in a FLUOstar Omega spectrophotometer (BMG Labtech, Ortenberg, Germany).

### qPCR

Liver samples were stored frozen at −80° before DNA extraction with the DNeasy blood and tissue kit (QIAgen, Crawley, UK) according to manufacturer’s instructions. The sequence was amplified using the following primers: 5’-TTCCGGGACTTTCGCTTTCC-3’ (sense) and 5’-CGACAACACCACGGAATTG-3’ (antisense). Amplification was detected and normalised against glyceraldehyde 3-phosphate dehydrogenase which was amplified using the following primers: 5’-ACGGCAAATTCAACGGCAC-3’ (sense) and 5’-TAGTGGGGTCTCGCTCCTGG-3’ (antisense). Amplification reactions were carried out using 5 μL of sample, 2.5 μmol/L of each primer, and SYBR green master mix using the Quantitect SYBR Green PCR Kit (QIAgen, Crawley, UK) for a 25 μL reaction. The amplification conditions were 95°C for 10 minutes followed by 40 cycles of 95°C for 15 seconds, 60°C for 1 minute, 72°C for 30 seconds. Data were processed with StepOne^TM^ software v 2.3 (ThermoFisher Scientific, Rockford, IL, USA).

### Statistics

Data were analysed =using GraphPad Prism 5.0 software, San Diego, CA, USA. Comparisons of continuous variables between two or more experimental groups were performed using the Student’s two-tailed *t* test or one-way ANOVA with Dunnett’s post-test for pairwise comparisons with WT or untreated *Asl*^*Neo/Neo*^ mice as indicated. *p* values <0.05 were considered statistically significant. For non-normally distributed data, a log transformation was used to compare groups. Figures show mean ± standard error of the mean (SEM). Kaplan-Meier survival curves were compared with the log-rank test.

### Study approval

All procedures were performed under UK Home Office licenses 70/6906 and 70/8030 and approved by institutional ethical review.

## ACKNOWLEDGEMENTS

We thank Mr I. Doykov, Dr F. Mazzacuva, Dr E. Reid, and Mr M. Wilson for technical assistance with LC-MS/MS and Mr B. Warburton, Mrs G. Sturges and Mrs S. Richards for technical assistance with animal work. This work was supported by Action Medical Research for Children Charity (grant GN2137). PBM is in receipt of a Great Ormond Street Hospital (GOSH) Children’s Charity Leadership award (V2516). PBM and PG are supported by the National Institute for Health Research Biomedical Research Centre at GOSH for Children NHS Foundation Trust and University College London. PG is a senior Welcome Trust fellow. SMKB and SNW received funding from ERC grant “Somabio” (260862). SNW received funding from MRC NC3Rs grant (NC/L001780/1). RK was part-funded by Borne charity. AAR and MPH receive funding from the UK Medical Research Council (MR/N026101/1). AAR is also funded by the EU Horizon 2020 (BATCure, 666918).

## AUTHOR CONTRIBUTIONS

JB, SH, SJH, SMB, PBM, PG, SNW contributed to overall study design. JB performed experiments, contributed to the overall study design and wrote the manuscript, which was edited by all co-authors. DPP, JH, ERF, JN, RK, NS, MH, MPH, BB, AR, AV provided substantial assistance in experiments. HP analysed plasma samples. DAR provided statistical assistance.

## CONFLICT OF INTEREST

The authors have no competing financial conflict of interest to declare.

